# Disruption of nuclear architecture as a cause of COVID-19 induced anosmia

**DOI:** 10.1101/2021.02.09.430314

**Authors:** Marianna Zazhytska, Albana Kodra, Daisy A. Hoagland, John F. Fullard, Hani Shayya, Arina Omer, Stuart Firestein, Qizhi Gong, Peter D. Canoll, James E. Goldman, Panos Roussos, Benjamin R. tenOever, Jonathan B. Overdevest, Stavros Lomvardas

## Abstract

Olfaction relies on a coordinated partnership between odorant flow and neuronal communication. Disruption in our ability to detect odors, or anosmia, has emerged as a hallmark symptom of infection with SARS-CoV-2, yet the mechanism behind this abrupt sensory deficit remains elusive. Here, using molecular evaluation of human olfactory epithelium (OE) from subjects succumbing to COVID-19 and a hamster model of SARS-CoV-2 infection, we discovered widespread downregulation of olfactory receptors (ORs) as well as key components of their signaling pathway. OR downregulation likely represents a non-cell autonomous effect, since SARS-CoV-2 detection in OSNs is extremely rare both in human and hamster OEs. A likely explanation for the reduction of OR transcription is the striking reorganization of nuclear architecture observed in the OSN lineage, which disrupts multi-chromosomal compartments regulating OR expression in humans and hamsters. Our experiments uncover a novel molecular mechanism by which a virus with a very selective tropism can elicit persistent transcriptional changes in cells that evade it, contributing to the severity of COVID-19.

## Main Text

Anosmia and hyposmia, complete and partial smell loss, respectively, constitute frequent and defining symptoms of SARS-CoV-2 infections ^1–5^. Although olfactory deficits are common among patients with upper respiratory infections, these symptoms are typically accompanied by rhinorrhea and nasal congestion that result in reduced airflow through the nasal cavity and physical insulation of olfactory sensory neuron (OSN) cilia from volatile chemicals. In contrast, anosmia in COVID-19 appears to be independent from conductive interference, as patients consistently deny associated nasal obstructive symptoms. These observations have prompted the acceptance of anosmia as a key symptom in identifying potential cases of SARS-CoV-2 infection^4^, especially in otherwise asymptomatic patients. Yet, the emerging link between SARS-CoV-2 infection and anosmia raises important mechanistic questions related to the molecular features of olfactory sensory neurons (OSNs). Specifically, OSNs do not express Ace2 and Tmprss2^6–8^, receptors essential for host cell entry by the virus. Thus, the virus may infect OSNs though a different set of extracellular host receptors, or the reported anosmia represents a non-cell autonomous effect^5,9^. Distinguishing between the two models has immense importance: the former implies that the olfactory epithelium (OE) could constitute a virus gateway to the CNS, through the OSN axons; while the latter could provide insight to the non-cell autonomous consequences of COVID-19 infection, currently considered the cause of severe complications and death.

To obtain a molecular understanding of the mechanisms contributing to COVID-19 induced anosmia, we performed RNA-seq analysis on olfactory epithelia obtained from 19 SARS-CoV-2 positive and 3 control subjects at time of death, in order to establish the transcriptional baseline of the human OE. These specimens were obtained from critically ill patients at the onset of the COVID-19 pandemic, prior to the groundswell support for the inclusion of smell and taste loss as essential symptoms. Thus, information on the concomitant presence of anosmia exists only for one patient who self-reported profound smell and taste loss at the onset of symptoms. Based on current projections of the prevalence of smell loss among COVID-19 patients, 40-70% of these subjects may have experienced abrupt olfactory deficits^5,10^. Without *a priori* knowledge of olfactory deficits linked to our specimens, we sought to identify the region of nasal mucosa most highly enriched for OSNs, focusing on the olfactory cleft, a region at the roof of the nasal cavity bridging the superior septum and middle turbinate bones (Extended Data Fig.1a). OE from this region contains high concentration of OSNs as demonstrated by detection of the mature OSN-specific olfactory marker protein (OMP) and the OSN-enriched LDB1 and confirmed by scRNA-seq (Extended Data Fig.1b, c). Thus, the analysis presented thereafter utilizes microdissections of this particular region of the nasal epithelium. The consistency of this approach in regards of OSN representation is demonstrated by plotting the levels of 30 OSN-specific markers, which were previously identified from mouse scRNA-seq experiments, across the 22 autopsies, revealing comparable representation between samples (Extended Data Fig. 1d).

Upon establishing conditions for extraction of human OE with consistent cellular representation, we asked if the viral RNA genome is detectable in infected samples by RNA-seq. Indeed, we detect the SARS-CoV-2 RNA genome in all the OEs from infected patients, but not in control OEs (Fig.1a). There is strong variability in the abundance of the viral genome between samples, with autopsies obtained less than 10 days after symptom manifestation demonstrating elevated viral loads at higher frequency (Fig.1b). To determine the identity of the infected cells, we performed fluorescent *in situ* RNA hybridization (RNA FISH) by RNA-scope using a probe for the negative strand of the Spike gene, which detects only the replicating virus. Labeling is sparse across the human OE and concentrated predominantly at the lamina propria and to a lesser degree at the most apical layer occupied by sustentacular cells (Fig.1c-e). Some fluorescent signal is detected at the OSN layer, however, there is no significant difference in the OSN staining patterns between control and SARS-CoV-2 infected OEs (Fig.1c-e), suggesting non-specific hybridization. Finally, it should be noted that the same RNA FISH probe used in infected lung sections revealed frequent cellular infection in this tissue (Extended Data Fig. 2). Thus, our data suggest that the virus predominantly replicates in non-neuronal cells of the nasal epithelium, consistent with the expression patterns of ACE2 and TMPRSS2^6–9^.

**Figure 1.**
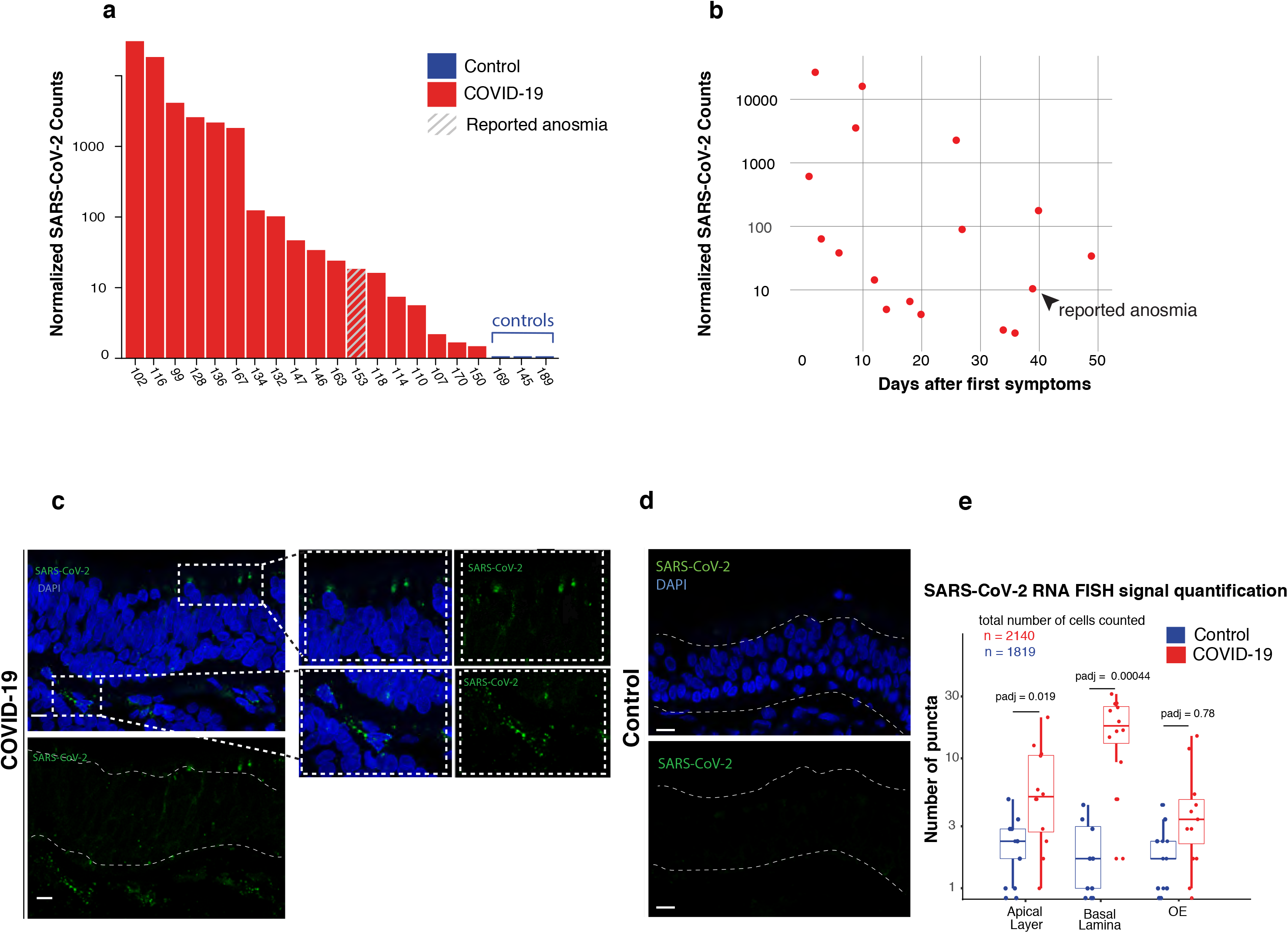
**a**, RNA-seq analysis of SARS-CoV-2 genomic reads in 19 human olfactory epithelium biopsies from COVID-19 patients (red) and 3 controls (blue). To account for differences in the viral load, SARS-CoV-2 raw counts were normalized to the hg38 genome and plotted as DESeq2’s median ratio normalization (MRN). The striped bar highlights the only sample with known anosmia. **b**, Normalized SARS-CoV-2 counts versus the number of days between the first symptoms of COVID-19 and autopsy collection. Arrow indicates the anosmic sample. **c,d,** Maximum intensity projections of confocal microscopy images of SARS-CoV-2 RNA FISH (green) in human olfactory epithelium of infected (c) and control (d) biopsies. This probe targets the antisense strand of the S gene, allowing detection of replicating virus. Nuclei are counterstained with DAPI. SARS-CoV-2 signal is detected in the apical layers of the epithelium, in close proximity to the sustentacular cells and basally, in the lamina propria. **e,** Quantification of RNA FISH signal at the apical, neuronal and basal layers of infected and control human OE sections suggests that the signal at the neuronal layer is non-specific whereas at the apical and basal layers is significantly enriched on infected vs control OE.

Since COVID-19 does not appear to infect OSNs, at least at a frequency that would account for the prevalence of olfactory deficits, we asked if the virus exerts its effects by influencing expression programs of specific cell types of the OE. Using previously described cell type specific markers (Extended Data Fig. 3a), we discovered that viral infection alters specifically the expression of mature OSNs (mOSNs) (Fig. 2a), the cell type responsible for detecting odors and for transmitting this information to the brain. Reduction of mOSN markers, however, does not stem from an increase in apoptotic markers in the infected OEs (Fig.2b). Moreover, not every mOSN marker is downregulated; genes with crucial role in OR signaling (Adcy3^11^, Cnga2^12^, Gng13^13^), and trafficking of ORs and Adcy3 (Rtp1^14^, Gfy^15^, respectively), are significantly downregulated in infected OEs (Fig, 2c, Extended Data Fig. 3a), whereas other mOSN-specific markers are not affected (Extended Data Fig. 3a). These observations prompted us to also explore the effect of SARS-CoV-2 infection in expression of the actual chemoreceptors *per se*. Strikingly, cumulative analysis of the total OR mRNA, shows that most infected OEs express significantly less OR mRNA than the three control samples (Fig. 2c, d). Infected OEs not only express less total OR mRNA, but also have lower complexity in the OR transcriptome, with 116 ORs being undetectable in most SARS-CoV-2^+^ samples (Fig. 2e). The age of the human subject does not appear to be a critical determinant of the expression properties of ORs and their signaling components (Extended Data Fig. 3b), refuting a bias in our analysis due to the increased death rates among older COVID-19 patients. Intriguingly, we do not detect a correlation between the total viral load at the human OE and the transcriptional effects on chemoreceptor pathways (Extended Data Fig. 3c), which is consistent with a non-cell autonomous process. In this vein, we asked if infected OEs exhibit excessive and prolonged pro-inflammatory responses, which have been reported systemically for patients with severe symptoms and could explain the reported olfactory deficits^16–18^. Surprisingly, we do not evidence for extreme or maladaptive anti-viral responses originating at the OE (Extended Data Fig. 3d), however we cannot exclude the systemic contribution of cytokines, or that such responses have been terminated in the OE by the time the patients succumbed. In support of the latter, the few samples collected within two days from the first symptoms tend to have elevated immune responses compared to samples with longer intervals (Extended Data Fig. 3e).

**Figure 2.**
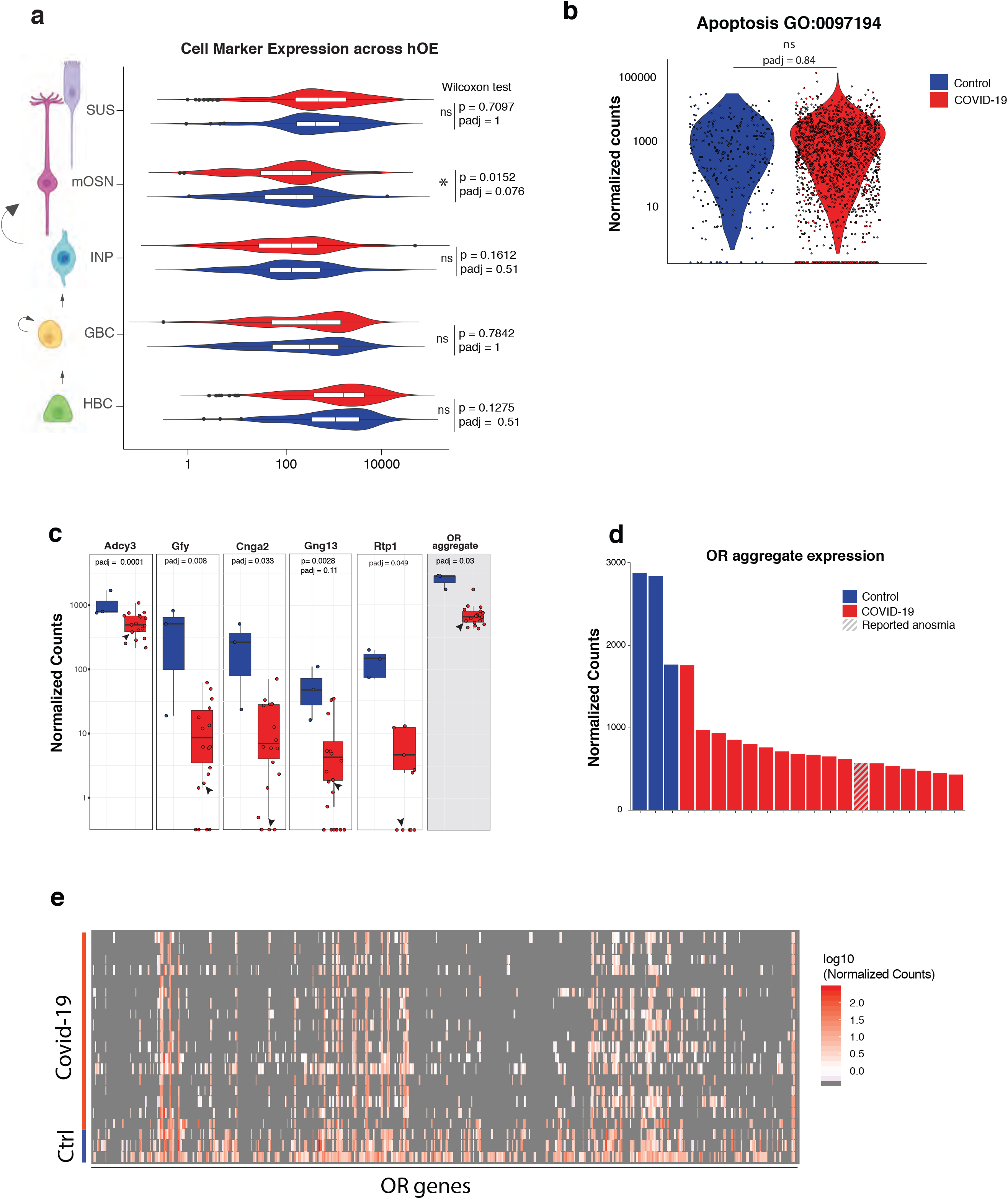
**a**, OE marker expression in SARS-CoV-2 infected and control specimens. Schematic representation of the cell types at different stages of differentiation, from the bottom: GBC (globose basal cells), HBCs (horizontal basal cells), INPs (immediate neuronal precursors), mOSNs (mature OSN) and SUS (sustentacular cells). A paired, two-tailed Wilcoxon rank sum test was used to determine whether the mean expression was different between SARS-CoV-2^+^ and controls. No significant changes were detected, except for mOSNs. **b,** Violin plot of apoptotic regulation response across all samples show no changes in distribution. p = 0.6891 was computed using Wilcoxon rank sum. The list of genes used corresponds to the GO category GO:0097194. **c,** Boxplot representation of the normalized counts (MRN) grouped in SARS-CoV-2^+^ positive and control samples for Adcy3, Gfy, Gng13, Rtp1, Cnga2, and aggregate OR mRNAs (highlighted in grey); *padj* values were generated with DESeq2 using the Benjamini–Hochberg method. Aggregate OR expression p value was calculated using two-tailed Wald test. Arrow indicates anosmic sample **d**, Distribution of aggregate OR mRNA across all human OE samples. The biopsy sample with known anosmia is highlighted with stripes. **e**, heatmap depicting expression of each OR gene in 3 control and 19 SARS-CoV-2 infected OE specimens.

To obtain independent confirmation for the SARS-CoV-2 induced downregulation of ORs and OR signaling molecules, and to explore early stages of infection, we performed experiments in golden hamsters (*M. auratus*). This rodent species is considered a good animal model for SARS-CoV-2 infection due to high sequence homology between hamster and human ACE2, and similarity in pathogenesis and immunological responses^19–22^. We performed a time course experiment that allowed us to monitor transcriptional changes in hamster OE, 1, 2, and 4 days after SARS-CoV-2 infection. As expected, we detect significantly higher viral loads in the hamster OEs compared to human epithelial specimens (Fig. 3a) due to the direct viral delivery into the hamster nasal cavities. Despite the high viral load, cellular infection in the hamster OE is infrequent, based on the immunoreactivity of the Nucleocapsid protein (NP) (Fig. 3b). Moreover, there is low correlation between NP and OMP immunoreactivity (Fig. 3c, Extended Data Figure 4a, b), consistent with our observations in human OE. Importantly, we do not detect NP in OSN axon bundles (Extended Data Fig. 4a, b) refuting the possibility that SARS-CoV-2 invades the CNS through the olfactory system. Consistent with the preferential non-neuronal tropism of SARS-CoV-2, we detect significant downregulation of sustentacular-specific markers from the earliest stages of infection (Fig.3d, Extended Data Fig. 4c). At later stages we also detect downregulation of markers specific for OSNs and their immediate precursors (Fig.3d, Extended Data Fig. 4c). Further, as previously reported^23^, we observe areas of restricted tissue damage (data not shown), coinciding with a weak, non-significant increase of apoptotic markers at 4dpi (Extended Data Fig. 4d). At this timepoint markers of the quiescent stem cells of the OE (Horizontal Basal Cells, HBCs) become upregulated (Extended Data Fig. 4c), suggesting that local tissue damage induced by high viral titers activates replenishment of Sus and OSN by HBCs. Reflecting the elevated viral titers at this acute infection timepoint, SARS-CoV-2 infection elicits strong inflammatory responses in hamster OE, beginning day 1 post infection (Fig. 3e). By day 4, there is a trend of dampening response, consistent with trends observed in human OE specimens. Importantly, there is strong and significant transcriptional downregulation of ORs and their signaling components, detected even by day 1 post infection (Fig. 3f, Extended Data Fig.4c). The kinetics and magnitude of transcriptional downregulation is not homogeneous across ORs; class I ORs, ~100 OR genes that are differentially regulated from the >1000 class II ORs, demonstrate slower and weaker downregulation than class II ORs (Fig. 3g). Notably, the same trend is observed in human OEs (Extended Data Fig.4e), suggesting that conserved differences in the regulation of these two OR subfamilies are responsible for the differential sensitivity to SARS-CoV-2 infection.

**Figure 3.**
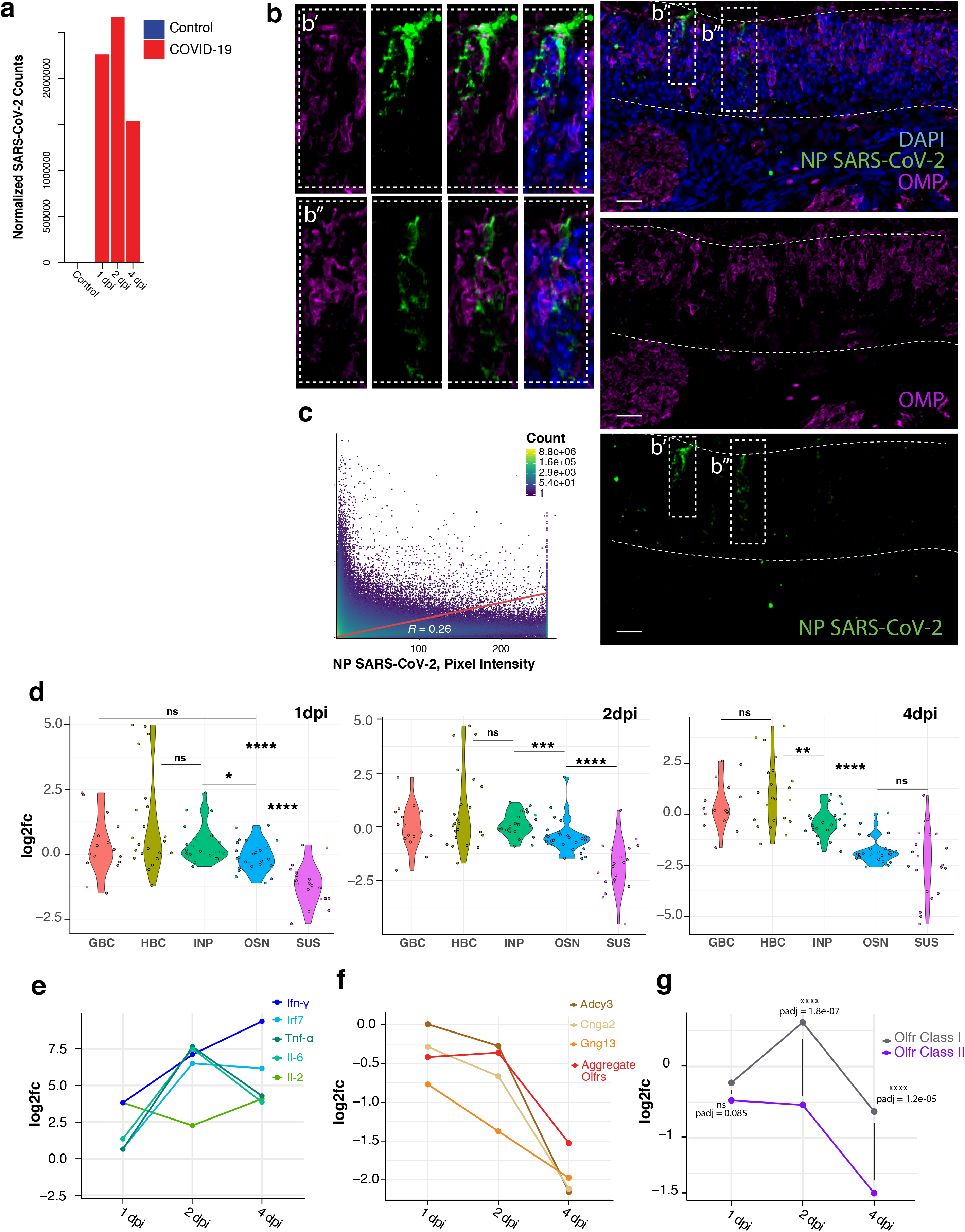
**a,** RNA-seq analysis of SARS-CoV-2 genomic reads in hamster olfactory epithelium following intranasal inoculation of SARS-CoV-2 and harvested at 1, 2, and 4 days post SARS-CoV-2 nasal infection (dpi). SARS-CoV-2 raw counts were normalized to the MesAur1.0 genome and plotted as DESeq2’s median ratio normalization (MRN). No mapped counts were found in the mock-injected control. **b,** Representative immunofluorescence image of co-staining against SARS-CoV-2 NP (green) and OMP (magenta). **c,** Colocalization analysis of OMP and SARS-CoV-2 NP shows no correlation between the two fluorescent signals. **d**, OE marker expression in hamster OEs 1, 2, and 4 days post SARS-CoV-2 infection, show immediate downregulation of sustentacular-specific markers, followed by delayed reduction of OSN and INP markers by day 4. **e**, Time course of inflammatory response genes Ifn-γ, TNF-α, Il-2, Il-6 plotted as log2FC at three different time points, 1dpi, 2dpi and 4dpi show increased expression of the inflammatory response. **f,** Time course plot of Adcy3, OR aggregate expression, Gng13 and Cnga2 log2FC, show consistent downregulation of ORs and OSN-specific genes involved in OR signaling. **g,** log2FC of class I and class II OR genes, reveals that class II ORs are significantly downregulated by day 2.

Because only class II OR gene expression depends on interchromosomal genomic compartments^24–26^, we asked if COVID-19 infection of the OE impacts the OSN nuclear architecture. To answer this, we established a protocol for the isolation of human and hamster OSN nuclei from crosslinked OE specimens by FACS (Extended Data Fig. 5a, b). *In situ* HiC on human OSN nuclei from two control and 4 SARS-CoV-2 infected OE samples revealed reduced genomic compartmentalization upon viral infection (Fig. 4a). Similarly, in hamsters, 3 days post SARS-CoV-2 inoculation (3dpi), there is a strong reduction in the number of predicted genomic compartments in OSN nuclei from infected OE, compared to controls (Fig. 4b). To increase the genomic resolution of our analysis towards identification of changes on specific compartments, we pooled the 2 control and 2 infected human OE specimens that are most distinct from each other. At a genome wide level, reduced genomic compartmentalization may be explained by an overall decrease on the frequency of specific genomic interactions and an increase of background interactions across all chromosomes, a result that is recapitulated in both humans and hamsters (Extended Data Fig. 5c, d). Crucially, we observe significant reduction in long-range *cis* and *trans* genomic interactions between OR gene clusters in both species upon SARS-CoV-2 infection (Fig. 4c-f). Quantification of genomewide *trans* genomic interactions composing OR-containing compartments, reveals strong and significant reduction in infected OEs from both species (Fig. 4g, h). Intriguingly, Adcy3, Gng13, and Gfy, together with Omp, Lhx2 and Atf5, which are also downregulated in infected human OEs form a separate interchromosomal compartment that also dissipates upon infection (Extended Data Fig. 6). Thus, the dramatic reorganization of nuclear architecture induced by SARS-CoV-2 infection does not only affect directly ORs, but also other genes that are essential for odor detection. Importantly, because we FAC-sorted nuclei with high levels of LHX2 and ATF5 proteins, disruption of genomic compartmentalization is likely the cause and not the consequence of Lhx2 and Atf5 downregulation, transcription factors that are critical for the expression of ORs and of their signaling molecules.

**Figure 4.**
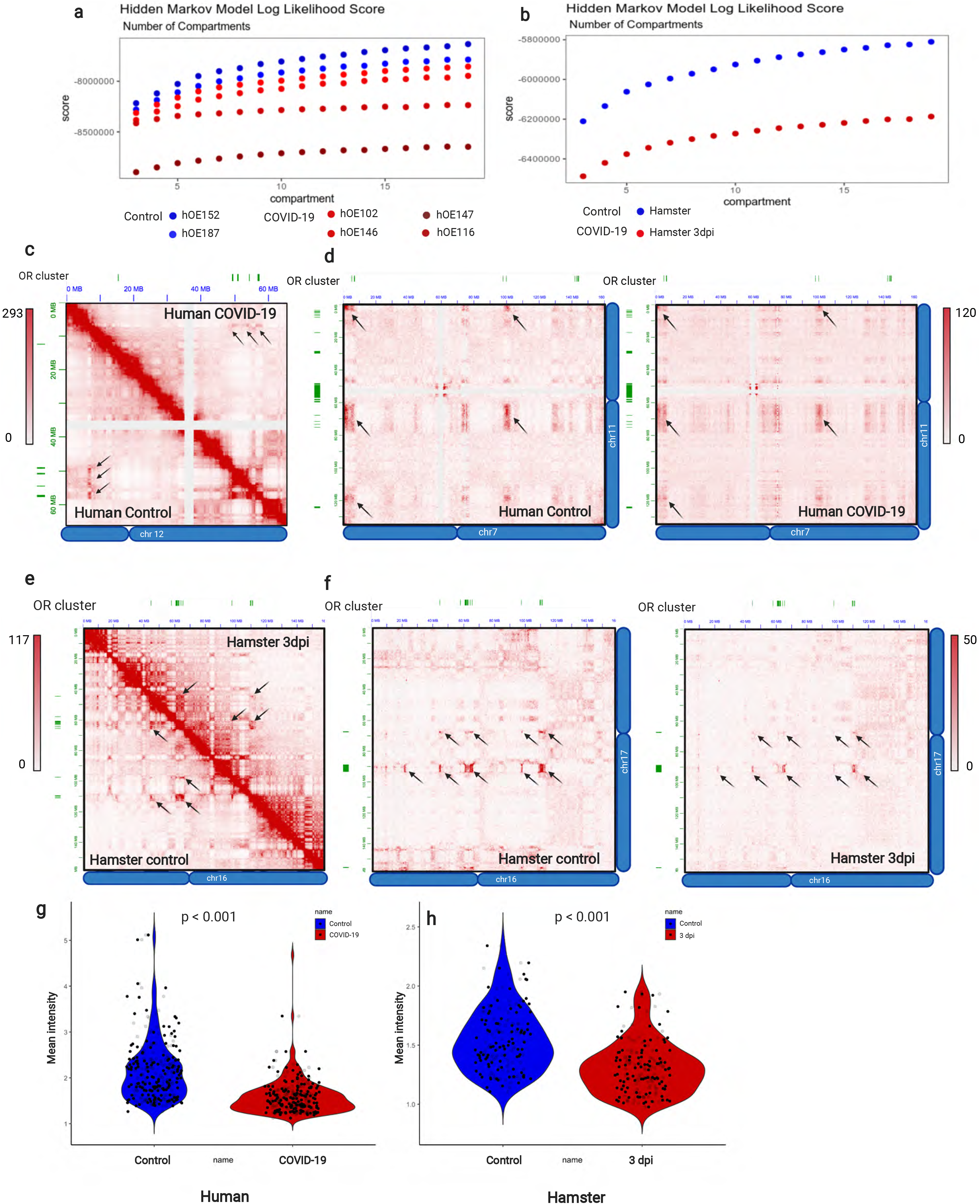
**a,** Machine-learning HMM score for a given number of compartments indicating different levels of genome compartments dissipation for control (blue) and COVID-19 patients (shades of red). For the rest of the panels, we pooled together human samples 152 and 187 (control) and 116 and 147 (SARS-CoV-2^+^) to increase the genomic resolution of our analysis and to identify changes occurring in the most disturbed nuclei. **b,** The same analysis for control (blue) and 3dpi infected hamster (2 control and 2 infected samples pooled together) (red). **c-f** Representative HiC maps of contacts between OR clusters in *cis* for human (**c**) and hamster (**e**) from pooled data. In each case control is the lower triangle below diagonal and SARS-CoV-2^+^ the upper triangle. Pixel intensity represents normalized number of contacts between pair of loci. Maximum intensity indicated at the top of each scale bar **d,f** Interchromosomal HiC contacts between OR clusters for human and hamster respectively. Genomic position of OR clusters indicated as green bars; arrows indicate the same OR compartments for both conditions. Pixel intensity represents normalized number of contacts between pair of loci. Maximum intensity indicated at the top of each scale bar. **g,h** Violin plot depicting the mean number of normalized *trans* HiC contacts between ORs from chromosome 11 to OR clusters genomewide at 50-kb resolution. Every dot indicates aggregated contacts between ORs on chromosome 11 to ORs from other chromosome, *p* value was computed using Wilcoxon rank test.

Our experiments reveal a molecular explanation for SARS-CoV-2 induced anosmia and uncover a novel mechanism by which this virus can alter the identity and function of host cells that lack entry receptors. Consistent with the previously reported absence of ACE2 and TMPRSS2 from OSNs, our data suggest that OSN infection by SARS-CoV-2 is too infrequent to account for the reported smell loss and the widespread downregulation of ORs and their signaling molecules. Although we cannot exclude the possibility that SARS-CoV-2 infects OSNs via interactions with Neuropilin-1^27^, our data do not support that viral replication in OSNs and transmission via their axons to the olfactory bulb^27,28^, as a cause of anosmia. The simplest explanation for the widespread olfactory deficits is a non-cell autonomous process, which may be mediated by the early onset induction of cytokines by the infected cells^29^. A direct consequence of this process is the dramatic reorganization of OSN nuclear architecture and dissipation of genomic compartments (Fig. 5). Nuclear reorganization deprives OSNs from the ability to detect odorants and to transmit this information to the brain, as both ORs and OR signaling molecules depend on these interchromosomal contacts for their expression. Given that most animals use olfaction for social communication, a virus induced loss of smell may limit social interactions of the infected individuals with their conspecifics^30^, limiting transmission within their social group. On the other hand, this process may reflect a viral effort to evade innate immunological responses since mammalian cells deploy interchromosomal contacts to activate antiviral programs^31^. Since adult CNS neurons also assemble long-range *cis* and *trans* genomic compartments^32,33^, this adaptive process may eventually induce long-lasting changes in brain nuclear architecture explaining cognitive and neurological deficiencies linked to SARS-CoV-2 infection^34^. Finally, it is worth noting that the realization that different OR sub-families exhibit differential sensitivity to SARS-CoV-2 infection, could eventually be deployed towards the development of rapid, COVID-19 specific smell-based screening tools.

**Figure 5.**
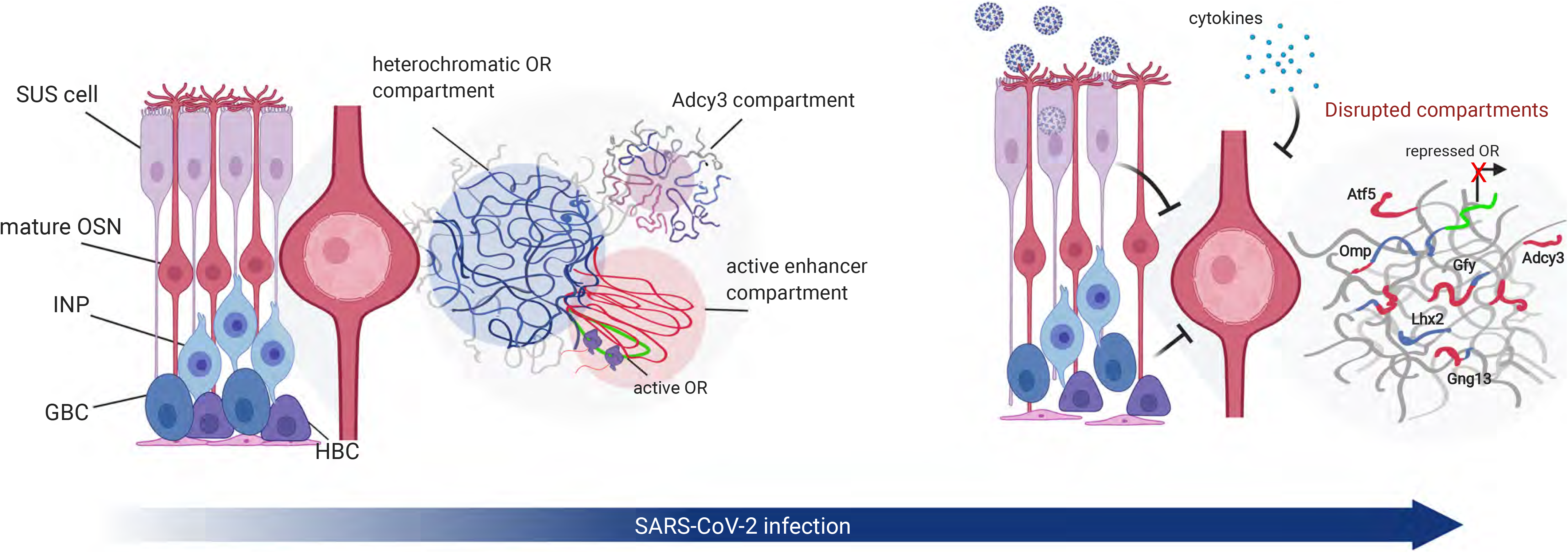
A model for the non-cell autonomous induction of anosmia by SARS-CoV2 infection. In control samples (left) silent ORs form a genomic compartment (blue) that promotes assembly of a multi-enhancer hub (red), which activates transcription of a single OR allele. Moreover, a second compartment consisted of numerous genes necessary for transcription, trafficking, and signaling of OR proteins is identified in control OSNs. Upon SARS-CoV2 infection (right), sustentacular cells or cells from the lamina propria elicit signals that induce disruption of genomic compartments and intermingling of genes that are supposed to be spatially segregated. Alternative, this disruption may be induced by elevated systemic cytokines circulating in COVID-19 patients. In either case, disruption of genomic compartmentalization results in downregulation of ORs and of their proteins involved in trafficking and OR signaling, resulting in anosmia.

## Acknowledgments

We thank members of the Lomvardas lab for critical comments and suggestions, Konstantin Popadin and Muhammad Saad Shamim for helpful analysis notes, David Weisz for assistance with software and Gary Struhl for the help with imaging, Charles Zuker, Tom Maniatis, Abbas Rizvi, and Max Gottesman for helpful comments and suggestions. The study was approved by the ethics and Institutional Review Board of Columbia University Medical Center (IRB AAAT0689, AAAS7370). LVG Golden Syrian hamsters (*Mesocricetus auratus*) were treated in compliance with the rules and regulations of IACUC under protocol number PROTO202000113-20-0743. This work was funded by 3R01DC018744-01S1 (NIDCD) (SL, JO) U01DA052783 (NIH Office of the Director, 4D Nucleome Consortium) (SL), HHMI Faculty Scholar Award (SL), and the Zegar Family Foundation (SL), R01AG067025 (P.R), R01AG065582 (P.R). Protocols and Sequencing data have been uploaded to the 4D Nucleome Data Portal (https://data.4dnucleome.org) and will be available to the public as soon as their curation is completed.

## METHODS

### Human samples

Patients previously diagnosed with COVID-19 by clinical symptoms and SARS-CoV-2 RT-PCR analysis when available, underwent full body autopsy at Columbia University Irving Medical Center (New York, NY, USA). The study was approved by the ethics and Institutional Review Board of Columbia University Medical Center (IRB AAAT0689, AAAS7370). **Hamsters:** LVG Golden Syrian hamsters (*Mesocricetus auratus*) were treated and euthanized in compliance with the rules and regulations of IACUC under protocol number PROTO202000113-20-0743. Detailed methods can be found in Extended Data. No statistical methods were used to determine sample size. The experiments were not randomized and investigators were not blinded to allocation during experiments.

## Extended Data

**Extended data Figure 1.**
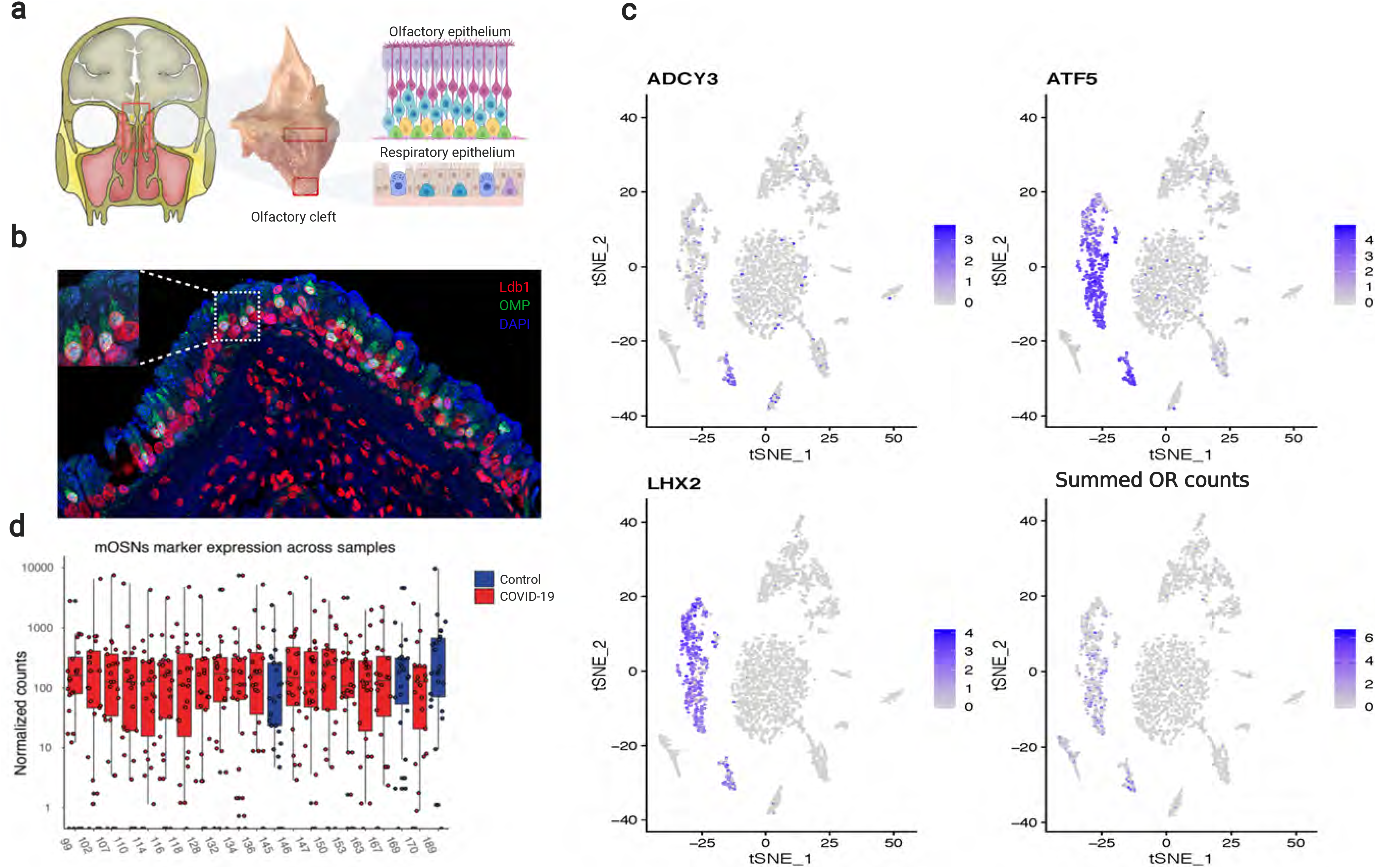
**a**, *En bloc* resection of the cribriform plate along with underlying mucosa from the olfactory cleft, which contains OE more superiorly and respiratory epithelium below. **b,** section of human olfactory epithelium stained for OMP (green) and Ldb1 (red), an OSN markers. Nuclei are labeled with DAPI (blue)**. c** tSNE plot representing clustering of FAC-sorted cell populations from the dissected region. ADCY3, ATF5 and LHX2 positive cells (blue) represent OSN identity. Summed OR counts confirm the cell identity. **d**, distribution of thirty OSN-specific markers previously identified from mouse scRNA-seq experiments across human samples used for the study.

**Extended data Figure 2.**
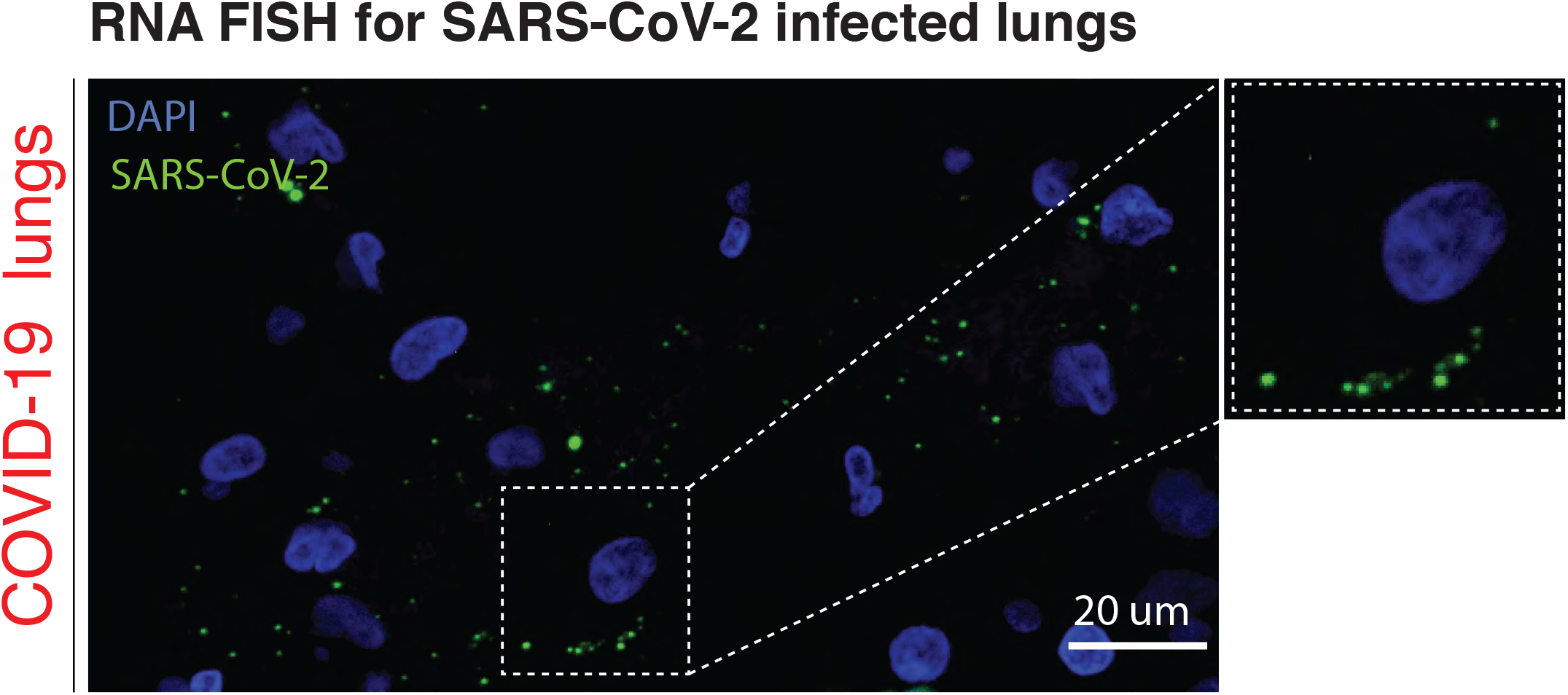
**a,** RNA FISH against SARS-CoV-2 in confirmed SARS-CoV-2^+^ human lung. Paraffin embedded lung sections show punctate fluorescent signal in proximity of nuclei counterstained with DAPI, showing the presence of abundant viral particles.

**Extended data Figure 3.**
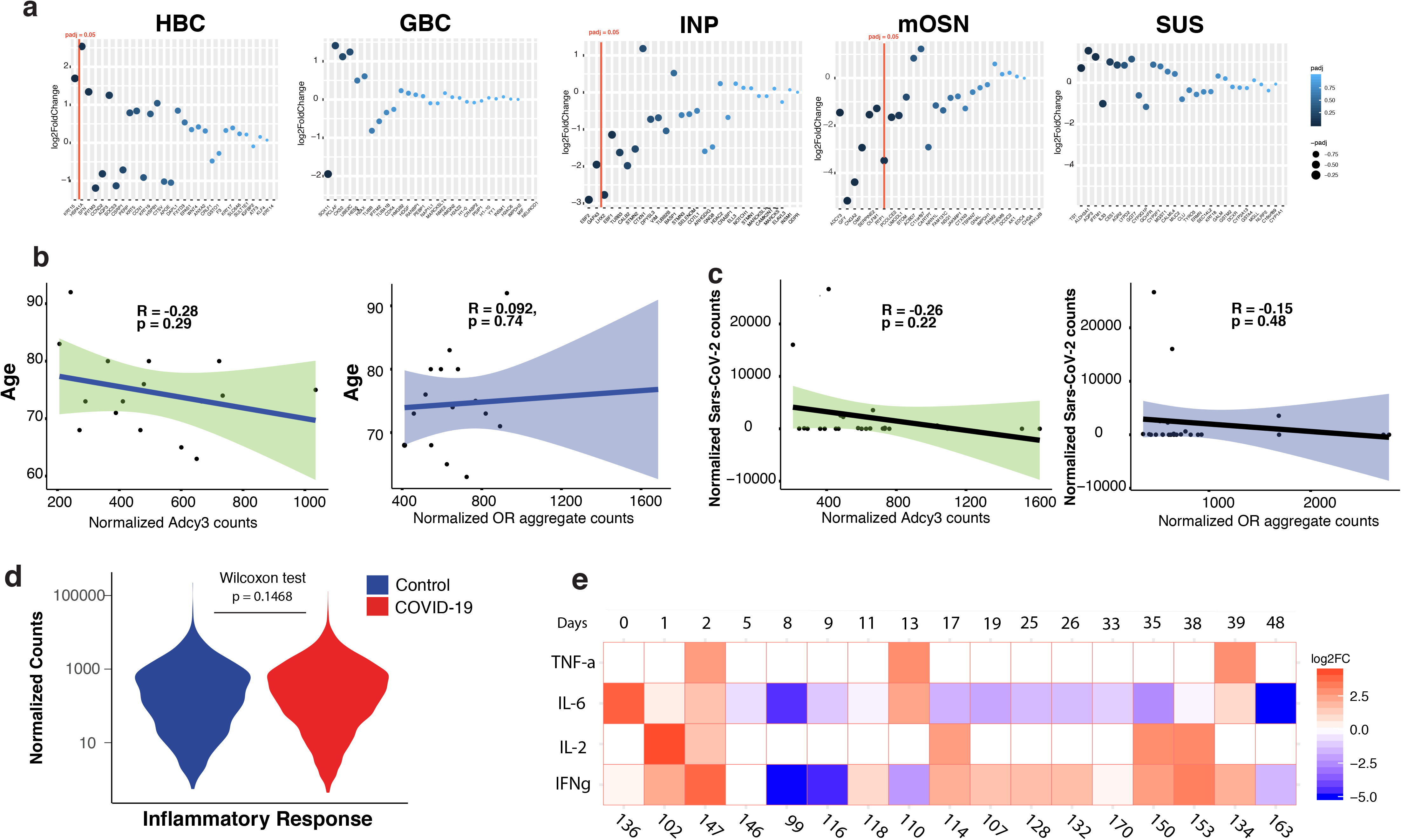
**a,** Log_2_ fold change expression of cell type-specific markers in the human OE. Red line depicts padj=0.05. Mature OSNs contain the highest number of significantly downregulated genes, followed by immediate neuronal precursors (INPs). **b,** Correlation of viral counts to normalized OR aggregate counts (green) and normalized Adcy3 counts (purple). The correlation coefficient and the significance level (p-value) were calculated with Pearson correlation test (p-value = 0.2699) and confirmed with Kendall’s rank correlation test (p-value = 0.2631) for Adcy3. Similarly, OR aggregate expression shows no significant correlation (p = 0.5319) which was confirmed with Kendall’s rank correlation test (p= 0.265). **c,** Regression analysis of Adcy3 and OR aggregate expression with age. No significant correlation was found in our cohort of samples which include individuals from 58 to 92 years of age. The statistical test used are the sample as reported in 3b. **d**, Violin plot of the inflammatory response across all samples show no changes in distribution. P = 0.1468 was computed using Wilcoxon rank sum. The list of genes used corresponds to the GO category GO:0097194. **e**, log2FC of Ifn-γ, TNF-α, Il-2, Il-6 on human OE specimens with different intervals from symptom manifestation to tissue harvesting (from 0 to 48 days).

**Extended data Figure 4.**
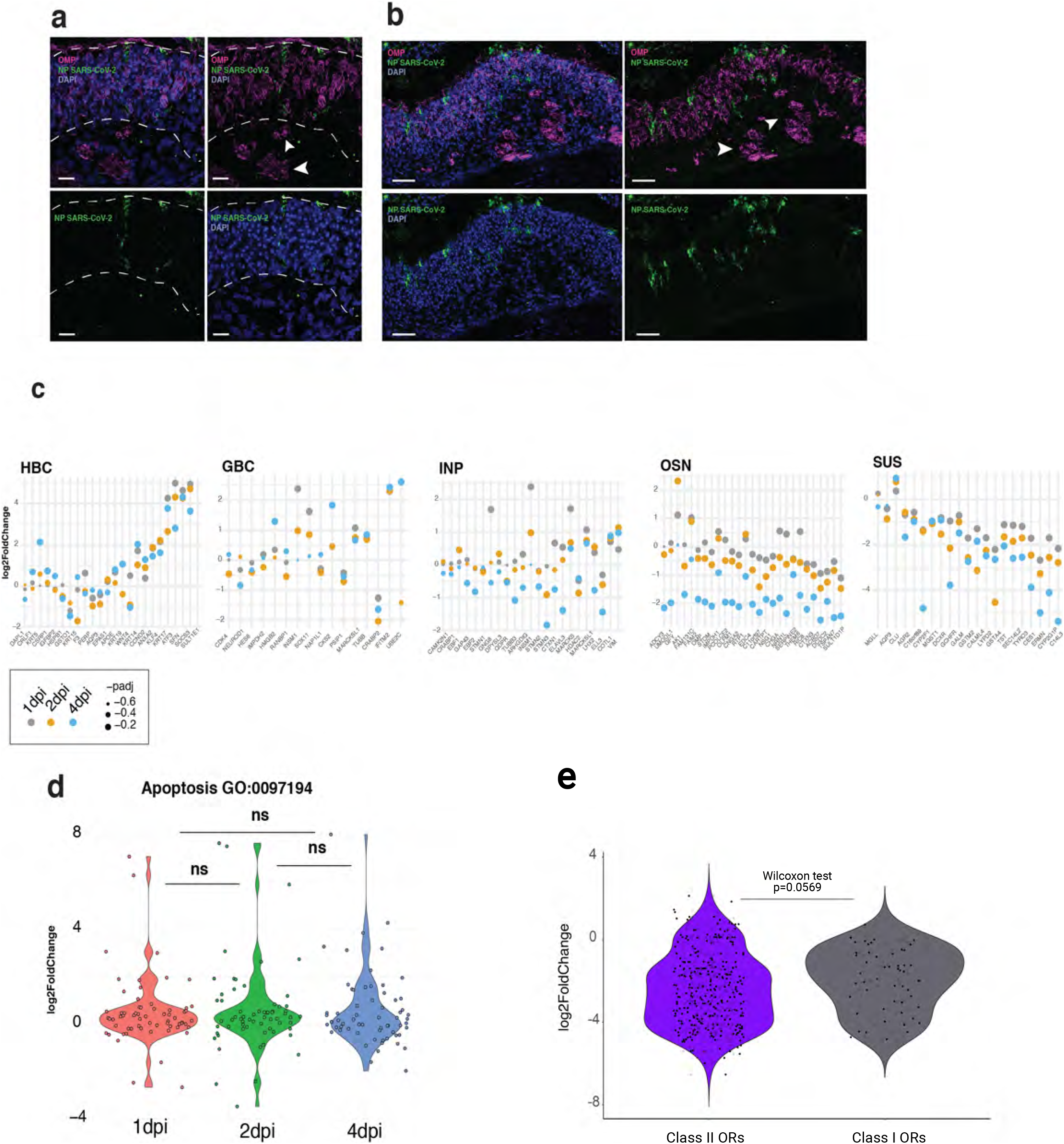
**a,** Immunohistochemistry for OMP (magenta) and SARS-CoV-2 nucleocapside, NP, (green) in hamster OE at 4dpi show one infected cell. White arrows indicate OMP-positive axon bundles which show no stain for SARS-CoV-2 NP. **b,** Immunohistochemistry for OMP (magenta) and SARS-CoV-2 (green) in hamster OE at 4 dpi. Representative image of an area of the tissue with multiple cells infected. No SARS-CoV-2 NP signal in the axon bundles was found. **c**, Log_2_ fold change expression of cell type-specific markers in hamster OE 1,2 and 3 days post infection. Sus markers are significantly downregulated from dpi 1, with mOSN and INP markers being downregulated at later days. **d**, Violin plot of apoptotic regulation response across all samples show no changes in distribution was computed using Wilcoxon rank sum. The list of genes used corresponds to the GO category GO:0042981. **e**, Log_2_ fold change expression of class I and class II ORs in human OE shows that that SARS-CoV-2 infection induces stronger downregulation of class II ORs.

**Extended data Figure 5.**
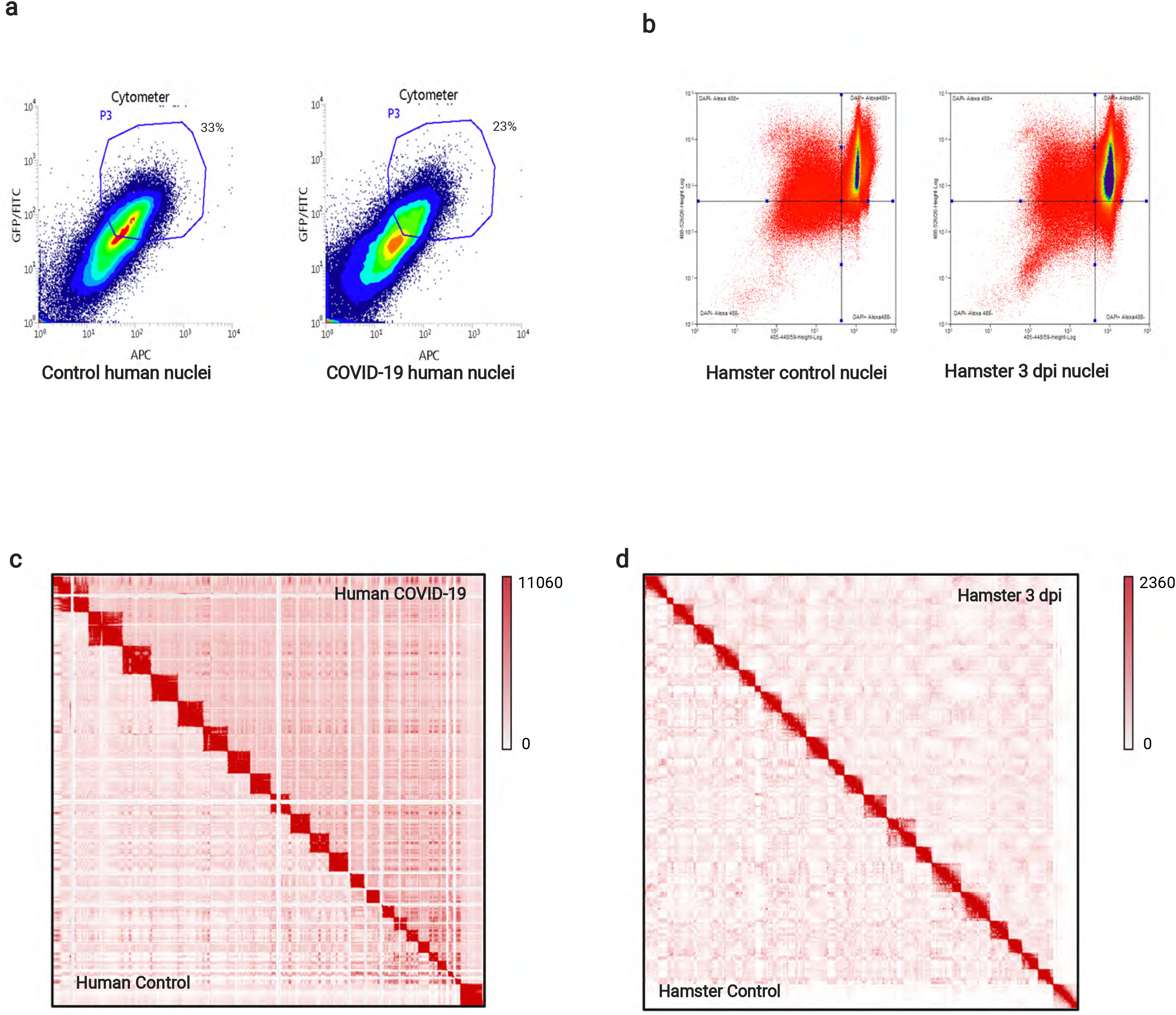
**a, b** Representative FACS data for control and SARS-CoV-2+ human and hamster specimens. Fixed DAPI positive, Lhx2/Atf5 double positive for human and Lhx2/OMP double positive for hamster, respectively, nuclei were collected for *in situ* HiC. **c,d** HiC map representing whole genome view on chromatin contacts for control (triangle below diagonal) and SARS-CoV-2 infected (triangle above diagonal) human and hamster respectively. **e,f** Chromatin compartments in human OSNs harboring Adcy3, Gng13 and Gfy in control (left **e,** top **f**) and SARS-CoV-2 samples (right **e**, bottom **f**). Arrows indicate compartments with genes involved (genomic annotation in green). **g,** annotated contact domains containing OR clusters (green bars); control domains propagate as blue ‘triangles’ below diagonal and for SARS-CoV-2 domains are indicated in ‘triangle’ above diagonal. **h,** the same for hamster.

**Extended data Figure 6.**
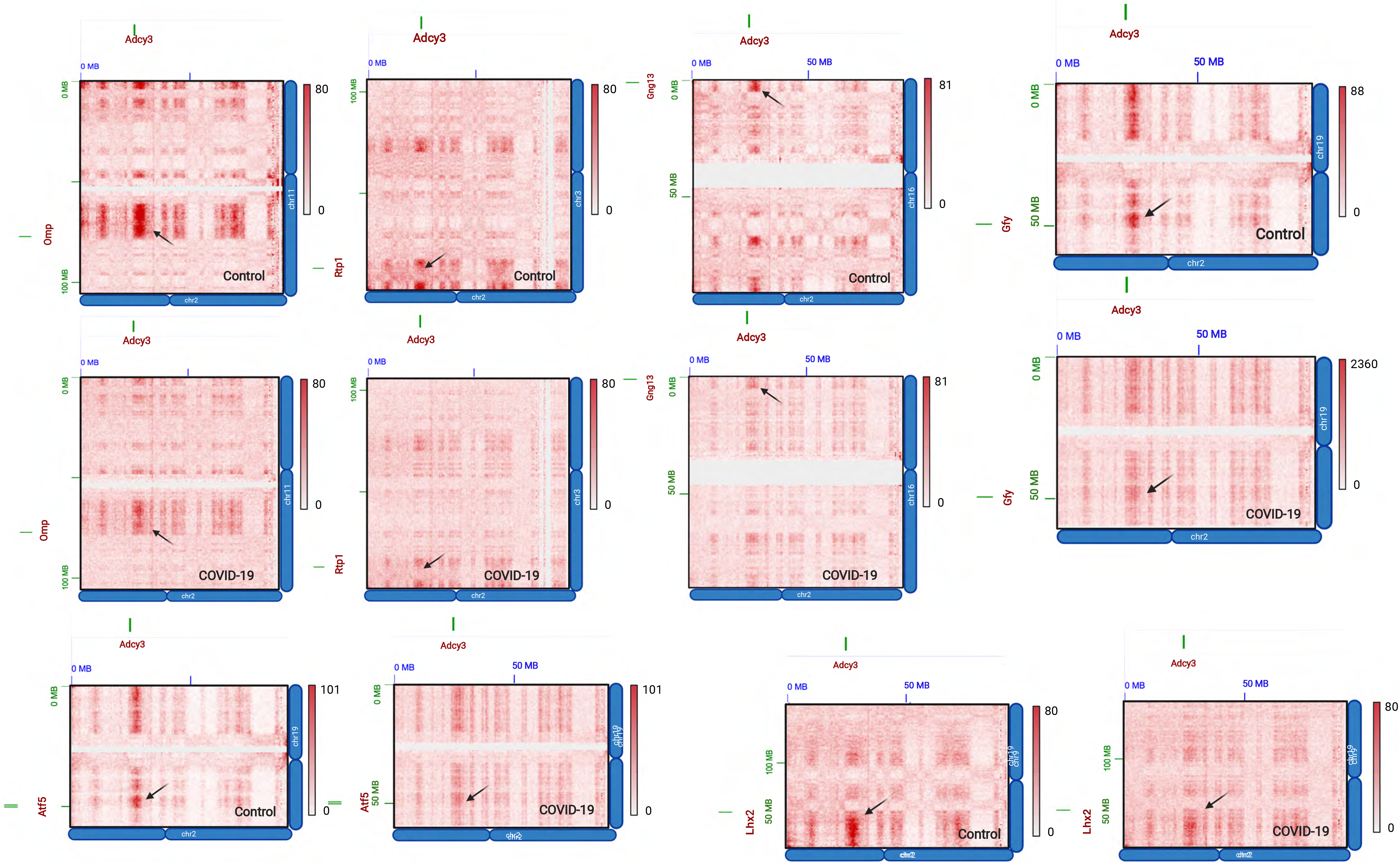
Chromatin compartments in human OSNs harboring Adcy3, Omp, Rtp1, Gng13, Gfy, Atf5 and Lhx2 in control (top and left in the bottom panel) and SARS-CoV-2 samples (bottom and right in the bottom panel). Arrows indicate compartments with genes involved (genomic annotation in green).

## Extended Data: Methods

### Human samples

Patients previously diagnosed with COVID-19 by clinical symptoms or SARS-CoV-2 RT-PCR analysis, underwent full body autopsy at Columbia University Irving Medical Center (New York, NY, USA) upon death. The study was approved by the ethics and Institutional Review Board of Columbia University Medical Center (IRB AAAT0689, AAAS7370). Specimens noted to have metastatic cancer and those exceeding a post-mortem time of 24 hours were excluded from subsequent analysis. Brain tissue and nasal epithelium, including the olfactory region, were retrieved under a collaborative effort by the Department of Neuropathology and the Department of Otolaryngology. Tissues were obtained and preserved for histological, molecular, and microscopic evaluation using separate surgical instruments to prevent cross-contamination. Control specimens were collected in similar fashion from deceased individuals who had no clinical history of COVID-19 and had negative SARS-CoV-2 PCR at the time of their presentation and again prior to post-mortem dissection. Nasal tissues, including olfactory and respiratory epithelium were harvested from the skull base using an en-bloc resection of the anterior skull base including the cribriform plate. Olfactory epithelium was isolated from the olfactory cleft, spanning turbinate and adjacent septal mucosa prior to being preserved in 1% paraformaldehyde (for HiC), 4% paraformaldehyde (for RNA ISH/IF), or Trizol (for RNA-seq).

### Hamsters

LVG Golden Syrian hamsters (*Mesocricetus auratus*) were treated and euthanized in compliance with the rules and regulations of IACUC under protocol number PROTO202000113-20-0743. Only male hamsters were used for experiments. All experiments were performed on dissected olfactory epithelium tissue or on dissociated cells prepared from whole olfactory epithelium tissue. Dissociated cells were prepared using papain (Worthington Biochemical) and FAC-sorted as previously described.

### Virus stock and propagation

Infectious work was performed at a CDC/USDA-approved BSL-3 facility at the Icahn School of Medicine at Mount Sinai. SARS-CoV-2 (clinical isolate UAS/WA1/2020) virus was propagated in Vero E6 cells in DMEM supplemented with 0.35% BSA. Infectious titer of virus was determined by plaque assay in Vero E6 cells using an overlay of Modified Eagle Medium (Gibco), 0.2%BSA (MP Biomedicals), 4mM L-glutamine (Gibco), 10mM HEPES (Fisher Scientific), 0.12% NaHCO_3_, 1% heat-inactivated FBS, and 0.7% Oxoid agar (Thermo Scientific). SARS-CoV-2 virus stocks used for hamster experiments were passage 3.

### SARS-CoV-2 inoculation

All hamster infections were performed in a BSL-3 animal facility at the Center for Comparative Medicine and Surgery at the Icahn School of Medicine at Mount Sinai (New York, NY) using 4-6-week-old male golden hamsters purchased from Charles River Laboratories. Hamsters were intraperitonially administered anesthesia of ketamine/xylazine (3:1), [100mg/kg] before inoculation. Inoculations were performed by intranasally administering 100 plaque-forming units (pfu) in a total volume of 100ul per hamster, diluted in PBS. Golden hamsters were provided thermal support after infection until recovery from anesthesia. Before sacrifice, the animals were anesthetized and then perfused with 60mL of PBS through the heart.

#### RNA-seq

RNA was extracted using Direct-zol RNA kits from Zymo Research. 50ng-1ug of total RNA was used to prepare DNA libraries with Truseq RNA Library Prep Kit v2 followed by 75 HO paired-end and multiplexed sequencing. Reads were aligned to human genome (hg38), *Mesocricetus auratus* (MesAur1.0) and SARS-CoV-2 (wuhCor1) using Subread and the raw read counts were assembled using featureCounts pipeline. Deseq2 was used to detect differences between conditions from the human samples and from the hamster biological replicates. For control samples we performed PCA analysis to remove outliers in an unsupervised fashion; two samples were not clustering with the rest and were removed, however Padj values for differences in OR, Adcy3, Gng13, Gfy and Cnga2 remain significant even if we keep these samples in our analysis (data not shown).

#### Single cell RNA-seq

Cells were dissociated according to the Worthington Papain Dissociation System by incubating fresh olfactory tissue with papain and Calcein VIolet for 40 min at 37 °C. Following dissociation, the live Calcein Violet-positive cells were sorted and assayed for scRNA-seq. Library preparation was performed accordingly to Chromium Single Cell 3ʹ v.3 Protocol and sequenced on NextSeq. Cell Ranger pipelines were used to generate fastq files which subsequently were aligned against hg38. All the cells with less than 1000 genes and less than 2000 UMIs per cell were discarded, resulting in 6828 cells used for analysis. Clustering was performed using Seurat with filtering default of a gene being expressed in at least 3 cells to include in data. To identify each of cell populations, previously annotated in lab olfactory epithelium markers were used to highlight olfactory lineages on tSNE plot.

#### RNAscope in hOE

Dissected tissue was fixed in freshly prepared 4% PFA for 24 hrs at 4C and soaked sequentially at 10%, 20% and 30% sucrose 1X PBS for cryopreservation. The tissue was embedded in OCT and 10 um thick sections were mounted on SUPERFROS Plus Gold slides. To detect the S gene transcripts of SarsCov2, RNAscope® Probe - **V-nCoV2019-S-sense**, cat# 845708, was incubated for 2 hr at 40C, in pre-treated sections as indicated by the RNAscope Multiplex Fluorescenct v2 Assay kit. Zeiss Zen2012 SP1 (v8.1.9.484) was used for capturing confocal images. Autofluorescence of the human OE sections was computationally removed using the ImageJ add-on function described in Baharlou et al^35^.

#### Immunofluorescence

Dissected tissue was fixed in freshly prepared 4% PFA for 24 hrs at 4C. OE was embedded in OCT and coronal cryosections were collected at a thickness of 12 μm. Antigen retrieval was performed with 0.01M citric acid buffer (pH 6.0) for 10 minutes at 99C. Sections were rinsed in PBS and after *permeabilization* with 1x PBS *0.1*% Triton X 100, slides were incubated in blocking solution (4% donkey serum +5% nonfat dry milk + 4% BSA + 0.1% Triton X-100) for 30 minutes at RT. Tissue sections were stained with primary antibodies against OMP^36^ (1:100 dilution) and NP (1:200 dilution, MyBiosource Cat. no.# MBS8574840). DNA was labelled with DAPI (2.5 μg/ml, Thermo Fisher Scientific Cat. no.# D3571). Primary antibodies were labelled with the following secondary antibodies: for OMP, anti-chicken IgG conjugated to Alexa-488 (2 μg/ml, Thermo Fisher Scientific Cat. no.# A-11055, RRID:AB_2534102), for NP, anti-rabbit IgG conjugated to Alexa-555 (2 μg/ml, Thermo Fisher Scientific Cat. no.# A-31572, RRID:AB_162543). Confocal images were collected with a Zeiss LSM 700 and image processing was carried out with Fiji (NIH).

### Fluorescence-activated nuclei sorting

Frozen 1% PFA-fixed tissue was mechanically crushed using Covaris Impactor and then nuclei were extracted with OptiPrep Density Gradient Medium according to the Sigma Millipore protocol. Following extraction and filtering two times through a 35-μm cell strainer, nuclei were stained with Lhx2/Atf5 antibodies for human samples and Lhx2/OMP antibodies for hamster respectively. Next DAPI/Lhx2/Atf5 and DAPI/Lhx2/OMP triple positive nuclei were sorted on a BD Aria II or BD Influx cell sorter for HiC experiments.

### In situ Hi-C

Depending on the sample, between 30 thousand and 100 thousand nuclei were used for in situ Hi-C. Sorted nuclei were lysed and processed through an in situ Hi-C protocol as previously described with a few modifications. In brief, cells were lysed with 10 mM Tris pH 8 0.2% Igepal, 10 mM NaCl. Pelleted intact nuclei were then resuspended in 0.5% SDS and incubated for 20 min at 62 °C for nuclear permeabilization. After being quenched with 1.1% Triton-X for 10 min at 37 °C, nuclei were digested with 25 U/μl MseI in 1× CutSmart buffer for 1.5 hours at 37 °C. Following digestion, the restriction enzyme was inactivated at 62 °C for 20 min. For the 45-min fill-in at 37 °C, biotinylated dUTP was used instead of dATP to increase ligation efficiency. Ligation was performed at 25 °C for 30 min with rotation after which nuclei were centrifuges. To degrade proteins and revers crosslinks pellets were incubated overnight at 75 °C with proteinase K. Each sample was transferred to Pre-Slit Snap-Cap glass mictoTUBE and sonicated on a Covaris S220 for 90 sec (‘Harsh shear’ program).

### Hi-C library preparation and sequencing

Sonicated DNA was purified with 2× Ampure beads following the standard protocol and eluted in 300 μl water. Biotinylated fragments were enriched as previously described using Dynabeads MyOne Strepavidin T1 beads. The biotinylated DNA fragments were prepared for next-generation sequencing directly on the beads by using the Nugen Ovation Ultralow kit protocol as described [Kevin/Adan paper]. DNA was amplified by 7 cycles of PCR. Beads were reclaimed and amplified unbiotinylated DNA fragments were purified with 1× Ampure beads. The quality and concentration of libraries were assessed using Agilent Bioanalyzer and Qubit Quantification Kit. Hi-C libraries were sequenced paired-end on NextSeq 500 (2 × 75 bp), or NovaSeq 6000 (2 × 50 bp).

### Hi-C data processing and analysis

Raw fastq files were processed using the Juicer single CPU BETA version on AWS. Human data were aligned against hg19 and hamster reads were aligned to MesAur1.0_HiC.fasta.gz using BWA 0.7.17 mem algorithm. Hamster genome assembly was obtained from the DNA Zoo Consortium^37^. After reads are aligned, merged, and sorted, chimaeras are de-duplicated and finally Hi-C contact matrices are generated by binning at various resolutions and matrix balancing. In this paper we present data with stringent cutoff of MAPQ >30. Hi-C matrices used in this paper were matrix-balanced using Juicer’s built-in Knight-Ruiz (KR) algorithm. Matrices were visualized using Juicebox^38^. Cumulative interchromosomal contacts at the 50 kb resolution were constructed by calling Juicer Tools dump to extract genome wide normalized data from a .hic file and subsequently analyzed as previously described^24^. A hidden Markov model (HMM) was used to assess the presence of genomic compartments as previously described^24^. Using 2–21 components, HMMs are constructed for odd versus even chromosomes and a score is calculated using hmmlearn’s built-in score to ascertain the likelihood of the given number of compartments. The same was done for even versus odd after transposing the matrix.

## Notes

### Competing Interest Statement

The authors have declared no competing interest.

